# Small molecule-mediated targeted protein degradation of voltage-gated sodium channels involved in pain

**DOI:** 10.1101/2025.01.21.634079

**Authors:** Alexander Chamessian, Maria Payne, Isabelle Gordon, Mingzhou Zhou, Robert Gereau

## Abstract

The voltage-gated sodium channels (VGSC) NaV1.8 and NaV1.7 (NaVs) have emerged as promising and high-value targets for the development of novel, non-addictive analgesics to combat the chronic pain epidemic. In recent years, many small molecule inhibitors against these channels have been developed. The recent successful clinical trial of VX-548, a NaV1.8-selective inhibitor, has spurred much interest in expanding the arsenal of subtype-selective voltage-gated sodium channel therapeutics. Toward that end, we sought to determine whether NaVs are amenable to targeted protein degradation with small molecule degraders, namely proteolysis-targeting chimeras (PROTACs) and molecular glues. Here, we report that degron-tagged NaVs are potently and rapidly degraded by small molecule degraders harnessing the E3 ubiquitin ligases cereblon (CRBN) and Von Hippel Lindau (VHL). Using LC/MS analysis, we demonstrate that PROTAC-mediated proximity between NaV1.8 and CRBN results in ubiquitination on the 2^nd^ intracellular loop, pointing toward a potential mechanism of action and demonstrating the ability of CRBN to recognize a VGSC as a neosubstrate. Our foundational findings are an important first step toward realizing the immense potential of NaV-targeting degrader analgesics to combat chronic pain.

## Introduction

Chronic pain is a major problem that affects more than 20% of Americans and has an economic burden of more than $600 billion annually^1^. This crisis led the NIH to launch the HEAL Initiative^2^, with one of the major goals being the identification of novel analgesics for the treatment of pain, with a focus on drugs that are non-addictive. Toward that goal, voltage-gated sodium channels (VGSCs) have emerged as attractive targets for non-opioid analgesia^3–5^. In particular, the channels NaV1.7 (*SCN9A*) and NaV1.8 (*SCN10A*), hereafter referred to as NaVs, have come under focus because of their preferential expression in the peripheral nervous system and the strong evidence implicating them in human pain^6–15^. Accordingly, there has been intensive effort in both industry and academic settings to develop selective inhibitors of these channels to serve as novel, non-addictive analgesics. The outcomes of such efforts have largely been disappointing^6,16^. A recent and notable exception is the first successful human clinical trial of VX-548, a novel NaV1.8-selective inhibitor from Vertex Pharmaceuticals, which was tested in a Phase II clinical trial showing for the first time that a NaV1.8-selective inhibitor could produce safe and efficacious analgesia in humans that was superior to placebo in acutely painful conditions (abdominoplasty and bunionectomy)^17^. This landmark study marks a major achievement in the search for novel and non-addictive analgesics and stands in contrast to the multitude of failures of other NaV subtype-selective inhibitors for pain, for instance those targeting NaV1.7, where either lack of efficacy or unacceptable adverse effects have halted progress^18,19^. With NaV1.8’s position as a bona fide human analgesic target confirmed, it now remains to capitalize on this knowledge by developing novel and superior therapeutics to modulate NaV1.8 in a variety of pain conditions. With respect to NaV1.7, successful and safe targeting in humans remains elusive, although the therapeutic potential is evident; the challenge will be to develop an approach that achieves efficacy and circumvents side effects such as anosmia, cardiotoxicity and autonomic disruption, which has recently come to the fore from several groups^20,21^

An emerging drug development approach that is ideally suited to circumvent these obstacles is targeted protein degradation (TPD). All existing pharmacological agents for VGSCs work via target occupancy to transiently *inhibit* function^22,23^. In contrast, TPD modulates the biological output of a protein of interest (POI) by reducing its *abundance*^24^. TPD is achieved by purposefully redirecting the native cellular protein degradation machinery to a POI, harnessing either the ubiquitin-proteasome system (UPS) or endolysosomal system^25^. A variety of pharmacological TPD strategies have been devised. The first and most developed effector class is proteolysis-targeting chimeras (PROTACs), which are heterobifunctional small molecules comprising a target-binding moiety (TBM) and an E3 ligase-binding moiety (EBM) joined by a linker. An effective PROTAC co-opts the UPS by inducing a ternary complex between itself, POI, and an E3 ligase, resulting in ubiquitination of the target and subsequent proteasome-mediated elimination. Monovalent degraders, also called molecular glues, also use a similar degradative mechanism of inducing proximity between a POI and E3 ligase^22^. Collectively, we refer to PROTACs and molecular glues as small molecule degraders, which contrasts them from the many other emerging formats of degrader therapeutics which use antibodies, nanobodies, small protein binders, peptides or aptamers as their underlying components.

TPD possesses many distinct advantages over conventional small molecule inhibition, including (1) a catalytic mechanism of action (MOA) permitting sub-stoichiometric dosing relative to target, (2) a more durable and sustained duration of effect, especially for proteins with slow resynthesis rates, (3) increased selectivity owing to requirement for a stable ternary complex, (4) the ability to engage a target protein potentially anywhere on its surface as opposed to only at a functional site (e.g. enzyme active site), and (5) the ability to eliminate the entirety of a protein’s functions (e.g. scaffolding roles) as opposed to just a single function^23–25^. These promising features, along with promise preliminary results from clinical trials of PROTACs^26–28^, have generated tremendous interest and effort to develop degrader therapeutics for a multitude of diseases, especially in the oncology and immunology space^27–30^. There has been comparatively less advancement of TPD for neuroscience applications, with some effort targeting proteins in the central nervous system^31^. These advantages of small molecule-mediated TPD are well-matched to the challenges of targeting NaVs in particular, which require specific and high levels of perturbation for biological effect. Moreover, the unique morphology of somatosensory neurons would make a TPD approach to NaV targeting especially efficacious and long-lasting, given that synthesis of NaVs occurs in the cell body up to 1 meter away from the their final required destination for physiological function and repopulation may take days to weeks^32^. However, to date, there are no published reports of small molecule-mediated TPD applied to ion channels such as NaVs, and there are no degrader therapeutics in clinical usage or development for pain.

Accordingly, in this study, we sought to determine whether small molecule-mediated TPD could be applied to NaVs. We demonstrate that both NaV1.8 and NaV1.7 are highly amenable to small molecule-mediated TPD *in vitro* using conditional degron tag (CDT) systems, and provide evidence that NaV1.8 can serve as a neosubstrate of CRBN based on LC/MS-based detection of site-specific ubiquitination. Furthermore, we generated candidate PROTACs based on non-specific sodium channel blockers (NSSCBs) and failed to observe the potent degradation we saw with our CDT experiments, highlighting the need to discover new intracellular binders of NaVs to realize the promise of small molecule-mediated TPD for analgesia.

## Results

### NaV1.8 is highly amenable to PROTAC-mediated intracellular targeted protein degradation (TPD)

We initiated our studies focusing on NaV1.8, given its recent confirmation as a bona fide human analgesic target in the VX-548 study^17^. To determine whether NaV1.8 is amenable to small molecule-mediated intracellular TPD with a PROTAC, we turned to conditional degron tag (CDT) systems, which enable one to render any protein of interest (POI) potentially degradable with tool degraders. We first applied the dTAG system, wherein a mutant FKBP12^F36V^ protein (hereafter referred to as dTAG) is appended to a POI at the N- or C-terminus, making the POI:dTAG chimera addressable by the tool PROTACs dTAG-13 (CRBN-recruiting) or dTAG V-1 (VHL-recruiting^33^ The dTAG system has been shown to be especially versatile among CDTs and generally produces the most robust degradation across protein classes ^34^

We created chimeras of human NaV1.8 (*SCN10A*) with the dTAG element and epitope tag V5 appended to either the C-terminal (hNaV1.8-dTAG) or N-terminal (dTAG-NaV1.8) **(Fig.1A)**. These chimeras were introduced through transient transfection into the the neuronal cell line ND7/23, which has been demonstrated to support native-like NaV1.8 surface expression and localization^35^. Cells were treated with CRBN-recruiting PROTAC dTAG-13 or vehicle (DMSO) for 24 hours at various concentrations, followed by lysis and measurement of abundance of dTAG-hNaV1.8 and hNaV1.8-dTAG by CEIA. Both chimeras were degraded by dTAG-13 in a dose-dependent manner, but the C-terminal tagged version (hNaV1.8-dTAG) showed greater degradation at all concentrations, and near complete elimination at 1000 nM **(Fig. 1B-D).** We confirmed the specificity of the dTAG-13-mediated degradation by also treating cells with a negative control compound dTAG-13-NEG, which showed no difference from DMSO vehicle control **(Fig. 1B).** Because of the superior degradation of the C-terminal dTAG chimera, we performed all subsequent experiments with hNaV1.8-dTAG and its derivatives. To more fully assess the dose-response relationship of dTAG-13-mediated degradation and to enable a more facile and scalable detection method for degradation, we developed a HiBiT-tagged variant (hNaV1.8-dTAG-HiBiT) that permits detection with plate-based HiBiT assays. We leveraged the multiplexing capability of this HiBiT assay to also test whether NaV1.8 is susceptible to PROTAC-mediated TPD via Von Hippel Lindau (VHL), which is the most frequently used E3 ligase for PROTAC development besides CRBN^25^. For this purpose, we treated cells with the VHL-recruiting dTAG-v1 PROTAC alongside dTAG-13. In this way, we observed a clear dose-response and nearly complete elimination of NaV1.8-dTAG by both dTAG-13 (D_max_ 95.1% at 1 µM) and dTAG-v1 (D_max_ 97% at 1 µM) at 24 hours **(Fig. 1E).**

**Figure 1.**
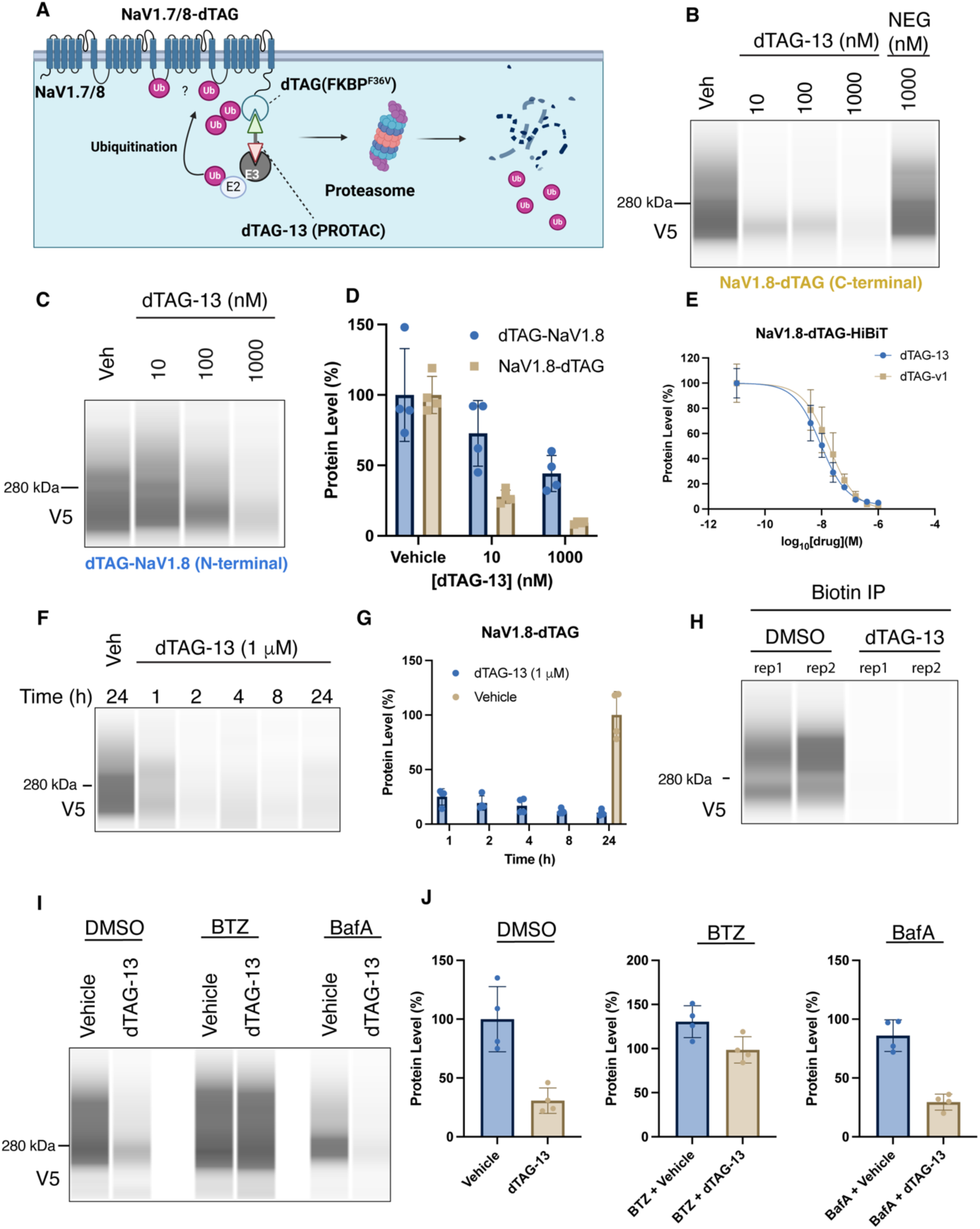
NaV1.8-dTAG is potently and rapidly degraded by dTAG PROTACs. **A.** Schematic showing NaV1.7 and NaV1.8 with FKBP12^F36V^ (dTAG) appended to the intracellular C-terminus. Engagement with the dTAG PROTACs dTAG-13 (CRBN-recruiting) or dTAG-v1 (VHL-recruiting) leads to ubiquitination and subsequent degradation by the proteasome. **B.** Representative CEIA analysis of C-terminal NaV1.8 dTAG chimera (NaV1.8-dTAG) expressed in ND7/23 cells and treated with dTAG-13, vehicle (DMSO) or dTAG-13-NEG (negative control) for 24 hours. NB: Anti-V5 antibody used in this and all other CEIA experiments. **C.** Representative CEIA analysis of N-terminal NaV1.8 dTAG chimera (dTAG-NaV1.8). **D.** Quantification of CEIA analysis comparing C-terminal NaV1.8-dTAG and N-terminal dTAG-NaV1.8 expressed in ND7/23 cells, performed simultaneously and separately from the experiments in 1B and 1C. Each measured V5 signal was normalized for each capillary by comparison to total protein signal. The normalized signal (area) for each technical replicate (n=4) was then divided by the average normalized signal for the vehicle condition and expressed as a percentage. The resulting values indicate the change in protein compared to the vehicle control condition. Bars represent the mean and error bars represent standard deviation (SD). **E.** Dose-response relationship of degradation of NaV1.8-dTAG-HiBiT by dTAG-13 and dTAG-v1 using the HiBiT DLR assay. Change from vehicle (DMSO) expressed as protein level (%), as for CEIA. Each point representing the mean of n=4 technical replicates and error bars indicating SD. **F.** Representative CEIA analysis showing time course NaV1.8-dTAG degradation by dTAG-13 (1 μM). **G.** Quantitation of CEIA time course data from 1F, n = 4 technical replicates, bars representing means with SD. **H.** Cell surface biotinylation and CEIA analysis showing complete degradation of surface-expressed NaV1.8-dTAG by dTAG-13 (1μM). N=2 technical replicates (rep) per condition. **I**. Representative CEIA analysis showing degradation of NaV1.8-dTAG by dTAG-13 (100 nM) in the presence of proteasome inhibitor bortezomib (BTZ, 500 nM) or lysosome inhibitor bafilomycin A (BafA, 100 nM). Cells were pre-treated for 1h with inhibitors followed by 4h of dTAG-13 or vehicle treatment in combination with the inhibitors. **J.** Quantitation of inhibitor experiment shown in 1I, with bars representing mean and error bars representing SD, n=4 technical replicates.

We performed additional experiments to assess the kinetics of dTAG-13-mediated degradation. Following transfection, we treated cells with dTAG-13 at various time points and observed strong degradation as early as 1h and maximal degradation at 8h (**Fig. 1F,G)** by CEIA.

The electrically functional fraction of NaV1.8 is that which is expressed at the cell surface, as opposed to that which is contained intracellularly. Therefore, we sought to determine whether dTAG-13-mediated degradation affected the surface expressed fraction of NaV1.8-dTAG. To do this, we transiently transfected NaV1.8-dTAG into ND7/23 cells and treated with either vehicle (DMSO) or dTAG-13 (1μM) for 24 hours and then performed surface biotinylation and immunoprecipitation with neutravidin beads, followed by CEIA probing against the V5 epitope tag. In this way, we demonstrated that (1) NaV1.8-dTAG is expressed on the cell surface and (2) dTAG-13 treatment completely eliminates that surface fraction **(Fig. 1H)**.

Ubiquitination of integral membrane proteins acts as a post-translational modification that prompts internalization and transit through the endolysosomal system^36^ However, to date, the dTAG system has largely been shown to cause TPD through the ubiquitin proteasome system (UPS)^33,37^. To determine which endogenous protein degradation system is operative with hNaV1.8-dTAG, we treated transfected cells with dTAG-13 in the presence of the proteasome inhibitor Bortezomib (BTZ) or the lysosome inhibitor Bafilomycin A (BafA) and observed nearly complete blockade of degradation by BTZ but no effect of BafA **(Fig. 1I)**.

### NaV1.8 is degradable by multiple conditional degron tag systems

Given the robust and nearly complete degradation of NaV1.8 with the dTAG system, we wanted to ensure that our observed outcomes are a general phenomenon and not a specific artifact of the dTAG system. The dTAG element possesses 16 lysines that could themselves be ubiquitinated by the dTAG-13-induced proximity of CRBN, thus rendering NaV1.8-dTAG artificially sensitive to UPS-dependent degradation in a way that the unmodified wild-type NaV1.8 may not be. To clarify this fundamental question, we used another recently developed CDT system that employs a minimal protein tag of 36 amino length (SD40) that was evolved to bind the monovalent molecular glue degrader PT-179. Binding of PT-179 to SD40 creates a protein interface for the docking of CRBN, thereby promoting the proximity-induced degradation of whatever POI bears the SD40 tag. The benefit of this complementary approach is that SD40 has just three lysines and is very small, thus favoring the scenario where ubiquitination of NaV1.8 rather than the SD40 tag drives degradation. As with dTAG, we appended SD40 to the C-terminal of hNaV1.8 in tandem with a HiBiT tag for detection (NaV1.8-SD40-HiBIT). Because SD40 was evolved to optimally interact with human CRBN, we expressed hNaV1.8-SD40-HiBIT in HEK293T cells, which are of human origin. Cells were treated for 24h with PT-179 at various concentrations and protein abundance was measured using the HiBiT lytic assay. We observed dose-dependent degradation of hNaV1.8-SD40-HiBIT, with a Dmax of 71% at (10 μM). **(Fig. 2A).**

**Figure 2.**
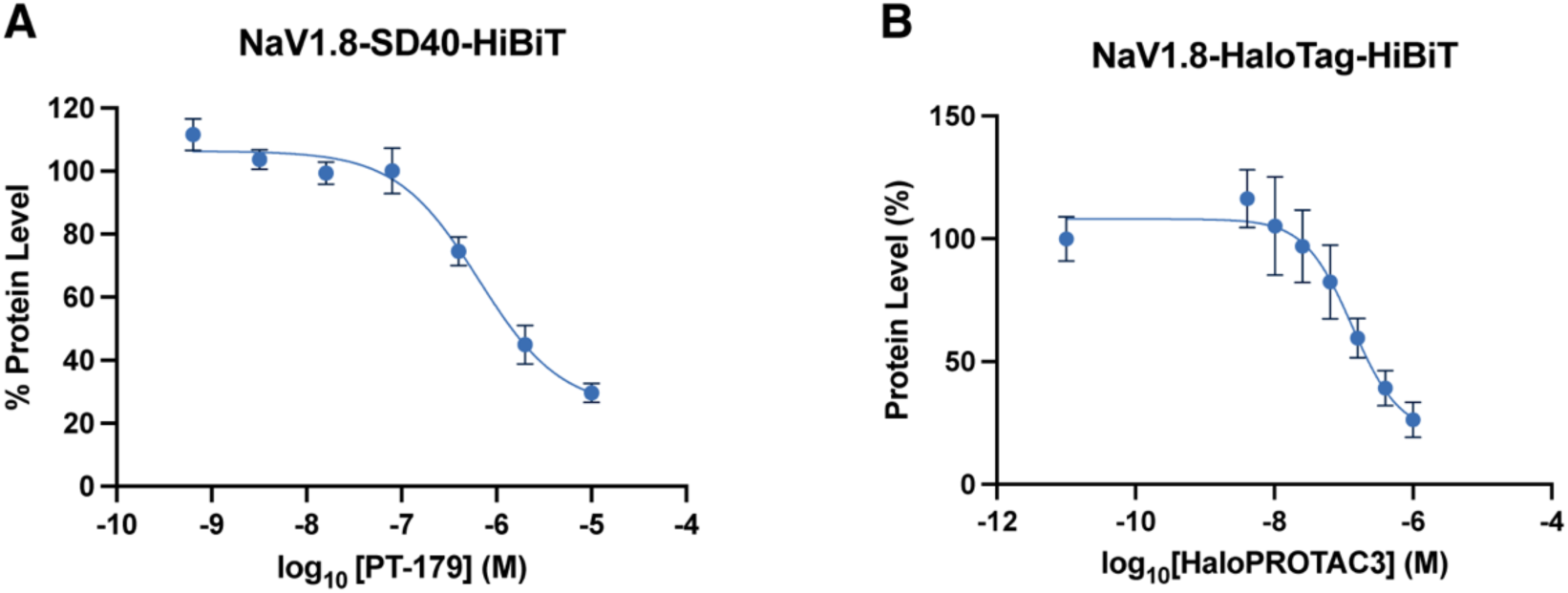
NaV1.8 is degradable with multiple conditional degron tag systems. **A.** NaV1.8-SD40-HiBiT dose-response curve with molecular glue PT-179 at 24 hours using the HiBiT DLR assay. Note: HEK293T cells used for this experiment, since SD40 was optimized for use with human CRBN. **B.** NaV1.8-HaloTag-HiBiT dose-response curve with HaloPROTAC3 at 24 hours using HiBiT DLR assay. Points represent means of n=4 technical replicates and error bars represent SD.

To further explore the generalizability of NaV1.8 targeted protein degradation, we developed a NaV1.8-HaloTag chimera to take advantage of the HaloPROTAC CDT system, which uses the HaloPROTAC3 compound to effect TPD via VHL^38^. Using HaloPROTAC3 at various concentrations for 24 hours, we observed strong degradation, achieving a D_max_ of 74% at 1 μM **(Fig. 2B)**. Neither SD40 nor HaloPROTAC systems achieved as thorough degradation as the dTAG system.

### PROTAC-mediated ubiquitination of NaV1.8-dTAG by CRBN occurs on the L2 intracellular loop

CRBN has emerged as a promiscuous E3 ligase that can recognize and ubiquitinate a diverse array of proteins that are not normally engaged by it, so called neosubstrates. VGSCs such as NaV1.8 have never been shown to be endogenous substrates of CRBN. Human NaV1.8 possess 63 lysine residues in its exposed cytoplasmic regions, all of which could be targets of CRBN. Therefore, to better understand the robust degradation we observe with dTAG-13, we aimed to identify the sites of ubiquitination on hNaV1.8-dTAG. To that end, we transfected ND7/23 cells with hNaV1.8-dTAG and treated them with dTAG-13 (1000 nM) or vehicle (DMSO) in the presence of BTZ to promote dTAG-13-mediated ubiquitination but halt degradation. Following treatment, cells were lysed and immunoprecipitation (IP) was performed using anti-V5 magnetic beads. The captured hNaV1.8-dTAG was digested on-bead and subjected to LC/MS analysis with data-independent analysis (DIA) to detect tryptic Lys-ɛ-Gly-Gly (diGLY) features, which are the remnants of ubiquitination and related ubiquitin-like proteins (e.g. neddylation)^39,40^. In this way, we detected a single significant enrichment of diGLY on K1100 (1.45 log_2_-fold dTAG-13/Veh), which resides in the 2^nd^ large intracellular loop (L2) of hNaV1.8 (**Fig. 3A,B**). Of note, no diGLY remnants were detected in the dTAG element.

**Figure 3.**
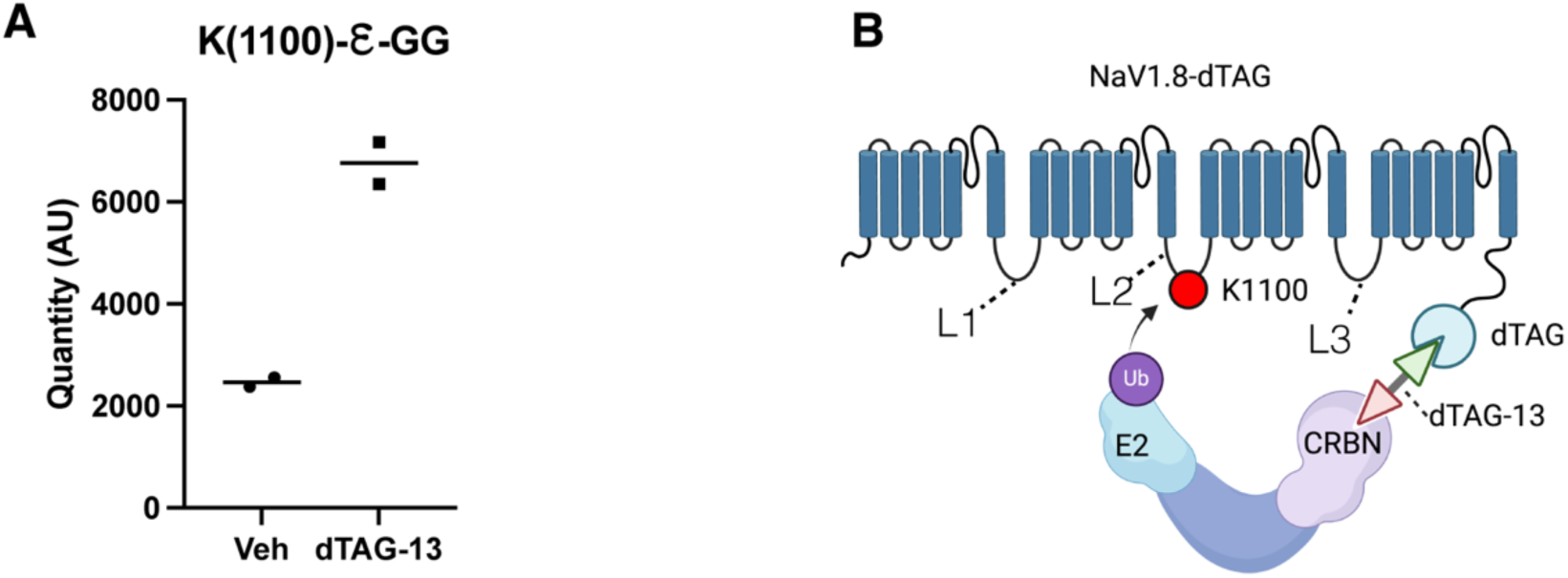
Chemically-induced proximity of CRBN and NaV1.8 by dTAG-13 results in ubiquitin modification on K1100 in the L2 intracellular loop. **A.** ND7/23 cells expressing NaV1.8-dTAG were treated with dTAG-13 or vehicle in the presence of BTZ for 3h, followed by anti-V5 immunoprecipitation, on-bead digestion and LC/MS analysis for tryptic diGly (K-ε-GG) remnants indicating a ubiquitin modification. dTAG-13 results in 1.45 log_2_-fold increased signal (arbitrary units, AU) compared to vehicle on K1100, which resides in the L2 intracellular loop region, depicted in **B.**

### PROTAC-mediated TPD extends to NaV1.7

Given the rapid and thorough degradation of hNaV1.8 by various methods, we wondered whether other high value ion channels are also amenable to PROTAC-mediated TPD. NaV1.7 is another VGSC that has been extensively studied and implicated in human pain, and thus represents a high priority target for analgesic discovery. Accordingly, we generated C-terminal (NaV1.7-dTAG) and N-terminal (dTAG-NaV1.7) dTAG chimeras with NaV1.7 to test its amenability to PROTAC-mediated TPD. NaV1.7 dTAG chimeras were transiently transfected into HEK293T cells and treated with a range of concentrations of dTAG-13 for 24 hours, followed by CEIA measurement of protein abundance. We observed strong dTAG-13-mediated degradation of the C-terminal NaV1.7-dTAG, with near complete elimination at 1 μM (**Fig. 4A,B**). Conversely, as with with NaV1.8, we observed only partial degradation of the N-terminal dTAG-NaV1.7, with a D_max_ of 51% at 100 nM and apparent hook effect at 1 μM (**Fig. 4C,D**). We also tested whether VHL could effect TPD of NaV1.7 using the VHL-recruiting degrader dTAG-v1. Treatment of transfected cells at 24 hours with dTAG-v1 resulted in nearly complete elimination of NaV1.7-dTAG at 100 nM (**Fig. 4F,G**).

**Figure 4.**
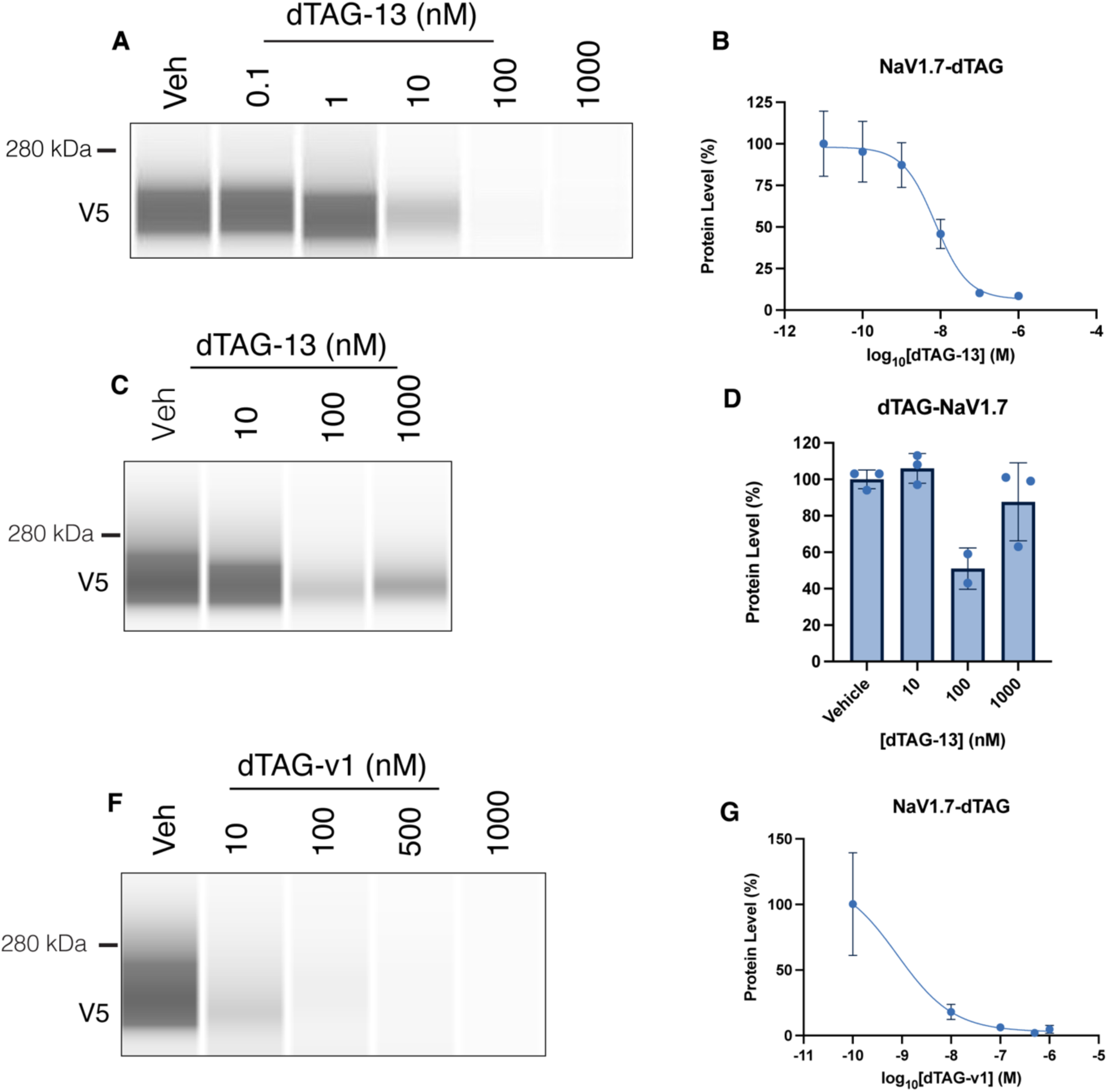
NaV1.7-dTAG is also highly degradable by dTAG PROTACs. **A.** Representative CEIA showing degradation dose response of C-terminal NaV1.7-dTAG to dTAG-13 at 24h. **B.** Quantitation of dose-response experiment in 4A. Points represent mean and error bars SD, n=4 technical replicates. **C.** Representative CEIA showing partial degradation dose response of N-terminal dTAG-NaV1.7 to dTAG-13 at 24h. **D.** Quantitation of dose response experiment 4C, bars representing means of n=2-3 technical replicates and error bars representing SD. **F.** Representative CEIA showing degradation dose response of C-terminal NaV1.7-dTAG to dTAG-v1 at 24h. **G.** Quantitation of data from 4F, points representing mean of n=4 technical replicates, and error bars representing SD.

## Discussion and Conclusions

We show here that the therapeutically important voltage-gated sodium channels NaV1.8 and NaV1.7 are amenable to small molecule-mediated TPD. To the best of our knowledge, our findings are the first demonstration that ion channels of any kind can be degraded via intracellular TPD with a small molecule PROTAC or molecular glue degrader. Using multiple conditional degron tag systems including dTAG, HaloTag and SD40, we show that both NaV1.7 and NaV1.8 are potently degraded, achieving rapid and near complete elimination *in vitro* with both CRBN and VHL-recruiting degraders. Further, we showed that these effects depend on the proteasome but not lysosome, and we provide evidence that PROTAC-induced proximity between CRBN and NaV1.8 results in ubiquitination in the L2 intracellular loop of NaV1.8, demonstrating for the first time that CRBN can recognize a VGSC as a neosubstrate. We attempted to develop prototype pan-NaV PROTACs using NSSCBs, which are the only class of small molecule known to bind to the intracellular part of NaVs. We did not observe any degradation of NaV1.8 with these compounds, indicating that NSSCBs are unlikely to be productive TBMs for TPD of NaVs and that new chemical matter will need to be developed to ultimately create effective NaV-targeting PROTACs.

Studies of TPD and PROTAC development have been heavily weighted toward oncology and immunology applications^25^, with some recent efforts in neuroscience, for example ARV-102 from Arvinas targeting Leucine-rich repeat kinase 2 (LRRK2) for Parkinson’s Disease. Our study stands as important step toward expanding the power and clinical potential of TPD into neuroscience, with a specific focus on the urgent and high impact problem of chronic pain.

The localization, trafficking and abundance of VGSCs and other ion channels are regulated endogenously by ubiquitination and deubiquitination^41^. A prominent and well-studied component of this endogenous regulatory network is NEDD4L (neural precursor cell expressed developmentally down-regulated 4-like), which is an HECT-family E3 ligase that ubiquitinates many types of membrane proteins at PY (PPxY) motifs, including NaV1.7 and NaV1.8^41^. An influential paper from Laedermann *et al.* showed that nerve injury downregulates NEDD4L in dorsal root ganglia (DRG) neurons, leading to a NEDD4L-dependent increase in surface expression of NaV1.7 and resultant neuronal hyperexcitability due to decreased endogenous ubiquitination^42^. Interestingly, they showed that ubiquitination by NEDD4L alters the surface localization of NaV1.7 but does not change the total abundance of this ion channel, indicating that NEDD4L-driven ubiquitination of NaV1.7 operates mostly in a non-degradative manner. The natural recognition of ion channels by NEDD4L has been exploited to manipulate their localization and abundance using protein- and peptide-based effectors. For example, a recent study by Sun *et al.* developed a genetically-encoded, virally-delivered chimeric protein comprised of the catalytic HECT domain of NEDD4L fused to a nanobody recognizing the β subunit of high-voltage-activated calcium channels, which they called Ca_V_aβaltor^43^. They directed this reagent toward the voltage-gated calcium channel Ca_V_2 in the DRG of mice and alleviated pain-like behaviors in the spared nerve injury model of neuropathic pain. Other studies from the same group (Colecraft) have demonstrated the ability to control the trafficking of many ion channels using the same general strategy of fusing POI-targeting nanobodies to ubiquitin-modulating effector components^44,45^. Recently, Martin *et al.* developed a lipidated peptide, PY(A), to promote endogenous NEDD4L-mediated degradation of NaV1.8 by disrupting the interaction between NaV1.8 and MAGI-1, a scaffolding protein that protects its binding partners from ubiquitination^46^. They used applied this PY(A) peptide *in vivo* and demonstrated pain reduction in the chronic constriction injury (CCI) model of neuropathic pain, and also showed functional reductions of NaV1.8 sodium currents in human DRG neurons *in vitro.* Taken together, these studies demonstrate the potential to harness the UPS to modulate NaVs for pain control.

Our findings stand in contrast to these prior studies in several ways. First, these prior approaches use protein or peptide-based effectors, while we used small molecule degraders, which can be fully chemically synthesized, do not require genetically-encoded expression, and are generally more stable and easier to administer in a therapeutic setting. Second, these prior works exploited the native recognition of NEDD4L for NaVs, while the small molecule degrader approaches used in our studies enabled non-native E3 ligase recruitment, in our case CRBN and VHL. NEDD4L-based effectors overall lead to changes in POI localization or trafficking but have more subtle effects on abundance, in some cases having no effect. In contrast, the CRBN- and VHL-recruiting small molecule degraders used here led to large reductions in NaV abundance, with near complete elimination of both NaV1.7 and NaV1.8 (D_max_ 95%) with dTAG-13 and dTAG-v1. These divergent outcomes likely arise from differences in the location, extent and type of ubiquitination caused by CRBN and VHL vs. NEDD4L, which warrants further exploration in future studies.

Our work has some limitations. We used multiple CDT systems, all of which demonstrated the amenability of NaVs for small molecule degrader-mediated TPD. Nevertheless, we cannot exclude that the degradation we observed was a consequence of ubiquitination of the degron tag and rather than the NaV. For example, the dTAG element possesses 16 lysines that could themselves be ubiquitinated, thus rendering NaV1.7/8-dTAG artificially sensitive to UPS-dependent degradation in a way that the unmodified wild-type NaV1.7/8 may not be. We partially addressed this possibility by using SD40, which is 36 amino acids in length and possesses only 3 lysines, thus favoring the scenario where ubiquitination of NaV1.8 rather than the SD40 tag drives degradation. Still, tag-driven ubiquitination and degradation cannot be entirely ruled out. In the absence of real NaV-targeting PROTACs, approaches that obviate the need for lysine-containing degron tags will be needed to resolve this question. For example, the recently reported method by Shade *et al.* using genetic code expansion to site-specifically attach E3 ligase ligands via click chemistry, would be useful for this purpose^47^. Another major limitation of our work is fact that we only evaluated degradation of NaVs *in vitro* with heterologous expression in immortalized cell lines. It remains to be determined whether and to what extent small molecule-mediated degradation of NaVs is possible *in vivo* and what the analgesic effects will be. Future studies using transgenic mice with degron tags knocked into the native NaV loci will be needed to address these questions, which we are currently developing. Our efforts to uncover to map sites of ubiquitin modification on NaV1.8 induced by dTAG-13 using LC/MS revealed lysine (K1100), but the diGly remnant could also indicate that neddylation or sumoylation was present at this site, not just ubiquitination. Moreover, our result does not permit one to conclude that ubiquitination at K1100 causes degradation, since the sensitivity of LC/MS analysis to detect diGly remnants without specific antibody-based enrichment is low, and our experimental approach likely left many potentially ubiquitinated lysines on NaV1.8 undetected. Indeed, there are 63 exposed lysines in the cytoplasmic region of NaV1.8, and it seems unlikely that only one bears any ubiquitin modifications. Future studies using di-Gly enrichment methods or tandem ubiquitin binding entities (TUBEs) will be needed to more accurately and specifically uncover the sites of ubiquitination that produce the robust degradation observed in our experiments.

In summary, our work demonstrates that NaV1.8 and NaV1.7 are amenable to small molecule-mediated TPD using PROTACs and molecular glues *in vitro* with CDTs. Due to the lack of intracellular binders of NaVs, it remains to be determined whether unmodified, wild-type NaVs can be degraded using a PROTAC or molecular glue. Answering this fundamental question and realizing the potential impact of small molecule-NaV-targeting degraders will require the discovery of novel intracellular small molecule binders of NaVs. Moreover, with the emergence of myriad extracellular TPD (eTPD) effectors, it will also be of great interest to understand whether NaVs and other ion channels can be degraded from outside the cell. Apart from therapeutic applications, our findings support the use of CDTs to control NaV abundance, which should find many uses in the study of these ion channels in preclinical pain neurobiology as a complement to genetic tools. Moreover, the approaches we demonstrate can serve as a precedent for TPD of other non-VGSC ion channels (e.g. PIEZO2, TRPV1) with broad application in neurobiology.

## Methods

### Reagents

dTAG-13 was obtained from Millipore Sigma and Tocris. dTAG-v1 and dTAG-13-NEG were obtained from Tocris. Bortezomib and Bafilomycin A were obtained from Cell Signaling Technologies.

### Construct design and cloning

All constructs were produced by VectorBuilder or Genscript using standard molecular cloning techniques. The base construct for all constructs in this study is a biscistronic mammalian expression vector with a human phosphoglycerate kinase (hPGK) promoter driving NaV transgene expression, and with a CMV promoter driving mCherry-Puromycin fusion reporter. Human NaV1.7 (Accession: NM_002977.3) and NaV1.8 (Accession: NM_006514.3) open reading frames (ORFs) were synthesized by Genscript and inserted into the base construct. Degron, epitope and HiBiT tags were then appended to the NaV ORFs. For C-terminal FKBP12^F36V^ (dTAG) and HaloTag constructs, the degron tag and adjoining linker elements were based on those from Bondeson *et al.*^34^, with the degron tag placed either at the C-terminal of NaV, separated by a rigid linker. The N-terminal dTAG-NaV1.8 and dTAG-NaV1.7 chimeras were made by appending FKBP12^F36V^ (dTAG) to the N-terminus via an SGLRSAT linker, as in Akin *et al*.^48^ For CEIA experiments, constructs included a V5 epitope tag, and for HiBiT experiments, a HiBiT tag was appended directly C-terminal to the dTAG element. SD40 sequences were obtained from Mercer *et al* ^49^ and appended directly C-terminal NaV1.8 without a linker. HiBiT tag was added to SD40 with a 3xGGGGS linker. All constructs were expressed in *E.coli* and delivered as transfection-grade purified supercoiled DNA. All constructs were quantitated using the Qubit dsDNA BR assay (Thermo Fisher Scientific; Q32850) prior to use. For HiBiT DLR assays, a construct in which firefly luciferase (FLuc) under the hPGK promoter was used.

### Cell Culture Experiments

ND7/23 cells were obtained from Millipore Sigma, and HEK293T cells were obtained from ATCC. Both cell lines were maintained using standard mammalian cell culture practices. Dulbecco’s Modified Eagle Medium (DMEM) supplemented with 10% Fetal Bovine Serum (FBS) and Penicillin-Streptomycin were used for both cell lines. For all experiments except NaV1.8-SD40-HiBiT, NaV1.8 constructs were expressed in ND7/23. NaV1.7 constructs were expressed in HEK293T. All transfections were performed using Lipofectamine 3000 (Thermo Fisher Scientific; L3000015) according to the manufacturer’s protocol. For plate-based assays (e.g. HiBiT), cells were initially seeded into T25 flasks for transfection, and then re-plated into 96- or 384-well assay plates for compound treatment and HiBIT assay. For CEIA experiments, cells were seeded into tissue culture vessels and transfected individually in their respective wells.

### Capillary Electrophoretic Immunoassay (CEIA)

Follow compound treatment, cells were lysed using Radioimmunoprecipitation assay (RIPA) buffer (Millipore Sigma; R0278) supplemented with cOmplete™, EDTA-free Protease Inhibitor Cocktail (Millipore Sigma; 11836170001). Lysates were quantitated using the Pierce BCA Protein Assay (Thermo Fisher Scientific; 23225) and subjected to the Simple Western 66-440 kDa Separation Module (SM-W008) according to the manufacturer’s instructions on the Wes or Jess instrument (Protein Simple). Primary antibodies used included rabbit monoclonal antibody against the V5 tag (Cell Signaling Technologies, 1320S) to detect dTAG constructs and rabbit monoclonal antibody against Vinculin (Cell Signaling Tecnologies, 13901S) as a load control. In some experiments, (Fig 1D,E), the Jess instrument was used and same-lane total protein normalization (instead of Vinculin) was used. Compass software (Protein Simple) was used to visualize and quantify the signal area for each sample, with load control or total protein normalization. Calculated corrected areas were analyzed in Microsoft Excel and GraphPad Prism (version 10).

### HiBiT Assays

The Nano-Glo® HiBiT Dual-Luciferase® Reporter System (Promega; CS1956A08) was used for all experiments, according to the manufacturer’s instructions. 96-well or 384-well white assay plates (Thermo Fisher Scientific; 165306 or 142762) were used, and luminescence measurements were performed using the BioTek Synergy H1 (Agilent). All data were exported from the Gen5 software (Agilent) into Microsoft Excel for initial data processing, which included background subtraction, per-well normalization to FLuc and then further normalization to the vehicle control condition (e.g. DMSO). Visualization and curve fitting was performed in GraphPad Prism using the log(inhibitor) vs. response -- Variable slope (four parameters) function.

### LC/MS Ubiquitin Site Mapping

Following transfection of NaV1.8-dTAG(V5) in 10 cm tissue culture plates, ND7/23 cells were pre-treated with bortezomib for 2 hours, followed by treatment with bortezomib and either DMSO or dTAG-13 for 2 hours. Cells were subsequently lysed in a 1% CHAPS immunoprecipitation buffer, 500 μl per plate, and lysates were incubated with ChromoTek V5-Trap® Magnetic Agarose (Proteintech; v5tma), 25 μl per samples, with end-over-end rotation at 4°C overnight. The following day, the beads were washed according to the manufacturer’s instructions. Captured protein was digested on-bead with trypsin using standard procedures at the Mass Spectrometry Technology Access Center at the McDonnell Genome Institute (MTAC@MGI) at Washington University School of Medicine. Purified peptides were run on the Vanquish Neo UHPLC System coupled to an Orbitrap Eclipse Tribrid Mass Spectrometer with FAIMS Pro Duo interface (Thermo Fisher Scientific). Data were acquired using the data-independent acquisition (DIA) method. Data were processed using Spectronaut 18 (Biognosys AG). The raw data files were analyzed via directDIA against the mouse SwissProt database, including the sequence of a custom protein (NaV 1.8), for GlyGly modification on lysine residue. Identification criteria were set to a false discovery rate (FDR) of 0.01 for peptide spectrum matches (PSMs), peptides, and protein groups, and a minimum post-translational modification (PTM) localization probability threshold of 0.75.

### Surface Biotinylation of NaV1.8

ND7/23 cells were transfected in 10 cm plates with NaV1.8-dTAG and then treated with either DMSO or dTAG-13 for 24 hours, after which cells were subjected to the Pierce Cell Biotinylation and Isolation Kit (Thermo Fisher Scientific; A44390) according to the manufacturer’s instructions, with the following modification. To avoid premature detachment from the plate, cells were labeled with NHS-biotin at 4°C, and after labeling, immediately transferred to a conical tube for further processing, which included neutravidin bead immunoprecipitation. Eluted protein was quantitated using the BCA assay as above and run on Simple Western 66-440 kDa size assay at a concentration of 2 mg/ml and probed with anti-V5 antibody, as above.

### Candidate NaV PROTAC synthesis

Compounds were synthesized at WuXi AppTec. Full synthetic methods can be found in Supplemental Material 1.

## Acknowledgements

This work was supported by departmental funds and a pilot grant from BioGenerator Ventures to AC, and by funds provided by the McDonnell Center for Cellular and Molecular Neurobiology at Washington University (RG). RG further acknowledges support of this work from the Dr. Seymour and Rose T. Brown Professorship in Anesthesiology. Mass Spectrometry analyses were performed by the Mass Spectrometry Technology Access Center at the McDonnell Genome Institute (MTAC@MGI) at Washington University School of Medicine, supported by the Diabetes Research Center/NIH grant P30 DK020579, Institute of Clinical and Translational Sciences/NCATS CTSA award UL1 TR002345, and Siteman Cancer Center/NCI CCSG grant P30 CA091842.

